# High-resolution mapping of Sigma Factor DNA Binding Sequences using Artificial Promoters, RNA Aptamers and Deep Sequencing

**DOI:** 10.1101/2024.10.18.619134

**Authors:** Essa Ahsan Khan, Christian Rückert-Reed, Gurvinder Singh Dahiya, Lisa Tietze, Maxime Fages-Lartaud, Tobias Busche, Jörn Kalinowski, Victoria Shingler, Rahmi Lale

**Affiliations:** Department of Biotechnology and Food Science, Faculty of Natural Sciences, Norwegian University of Science and Technology, Trondheim, Norway; Syngens AS, Trondheim, Norway; Bielefeld University, Center for Biotechnology (CeBiTec), Technology Platform Genomics, Bielefeld, Germany; Bielefeld University, Medical School OWL, Bielefeld, Germany; Department of Molecular Biology, Umeå University, Umeå, Sweden

**Keywords:** Sigma factor, Artificial promoters, RNA aptamers, *in vitro* transcription, Deep sequencing

## Abstract

The variable sigma (σ) subunit of the bacterial RNA polymerase holoenzyme determines promoter specificity and facilitate open complex formation during transcription initiation. Understanding σ-factor binding sequences is therefore crucial for deciphering bacterial gene regulation. Here, we present a data-driven high-throughput approach that utilizes an extensive library of 1.54 million DNA templates providing artificial promoters and 5^*′*^ UTR sequences for σ-factor DNA binding motif discovery. This method combines the generation of extensive DNA libraries, *in vitro* transcription, RNA aptamer selection, and deep DNA and RNA sequencing. It allows direct assessment of promoter activity, identification of transcription start sites, and quantification of promoter strength based on mRNA production levels. We applied this approach to map σ^54^ DNA binding sequences in *Pseudomonas putida*. Deep sequencing of the enriched RNA pool revealed 64,966 distinct σ^54^ binding motifs, significantly expanding the known repertoire. This data-driven approach surpasses traditional methods by directly evaluating promoter function and avoiding selection bias based solely on binding affinity. This comprehensive dataset enhances our understanding of σ-factor binding sequences and their regulatory roles, opening avenues for new research in biology and biotechnology.

---

Determining σ-factor binding sequences is critical for deciphering bacterial gene regulation. Despite the rapid expansion of sequenced genomes, our knowledge of how genes are regulated at the primary DNA sequence level lags behind. Even in well-studied model organisms like *Escherichia coli*, the regulation of over half the genes remains unclear^1^. This lack of understanding extends to other organisms, highlighting the need for a genome-wide approach to comprehend promoter regulation across all domains of life.

Transcription initiation, the first step of gene expression, is tightly controlled by σ-factors. In bacteria, RNA polymerase (RNAP) relies on σ-factors to recognize specific promoter sequences and initiate transcription. Different σ-factors exhibit distinct binding preferences, allowing bacteria to fine-tune gene expression in response to environmental cues^2,3^. However, comprehensive identification of σ-factor binding sequences remains a challenge. This knowledge gap hinders our ability to accurately predict and annotate promoter sequences, ultimately limiting our understanding of bacterial transcriptional regulation.

While traditional methods like DNA footprinting, Electrophoretic Mobility Shift Assays (EMSAs), Chromatin Immunoprecipitation (ChIP) coupled with next-generation sequencing (NGS), and Systematic Evolution of Ligands by Exponential Enrichment (SELEX) have provided valuable insights into σ-factor binding preferences, they have limitations hindering comprehensive identification^4–7^. These methods either rely solely on binding affinity or lack the resolution to specifically target σ-factor binding sequences within promoters. Additionally, pre-defined sequence libraries in protein-binding microarrays might miss important motifs, and SELEX prioritizes high-affinity binding, which might not be ideal for σ-factors where weak interactions still play an important biological role.

Our data-driven approach presented in this study overcomes many of these limitations by combining a comprehensive library of artificial promoters and the strengths of *in vitro* transcription (IVT) assays. This approach offers several key advantages. Primarily, we do a direct assessment of promoter activity through IVT reactions rather than relying solely on binding affinity. Additionally, by analyzing the enriched RNA pool, we pinpoint σ-factor binding sequences that initiate actual transcription, thus identifying functional binding sequences. Moreover, the use of an extensive DNA library and deep sequencing facilitates the discovery of a broad spectrum of σ-factor binding motifs, complete with quantitative data. This comprehensive data enables us to establish a genotype-phenotype linkage. Another significant benefit is the breadth of the data span. While identifying functional sequences, our approach also produces a substantial volume of non-functional DNA sequences. These non-functional sequences provide a crucial negative dataset for model training, which is essential for developing transcriptional models using machine learning. By incorporating both functional and non-functional sequences, we provide data allowing the machine learning models to distinguish between active and inactive promoter regions, ultimately improving their effectiveness in “understanding” transcriptional regulation.

To showcase the effectiveness of this approach, we applied it to the Gram-negative bacterium *Pseudomonas putida*. The experimental efforts reported in this study led to identification of 64,966 distinct σ^54^ DNA binding sequences, significantly expanding the known repertoire of DNA binding motifs for this particular σ-factor. This high-resolution data offers a multifaceted advantage. It not only deepens our understanding of σ^54^-dependent promoters in *P. putida*, but also lays the groundwork for developing powerful tools in synthetic biology, among others for rational engineering of promoters, training machine learning models for promoter prediction. Overall, this comprehensive and data-driven approach surpasses traditional techniques for identifying σ-factor binding sequences, paving the way for a deeper understanding of bacterial gene regulation.

## Results

### Construction of an Extensive Plasmid DNA Library with Diverse Regulatory Sequences

A critical component of our approach is the generation of extensive DNA libraries containing diverse 5^*′*^ regulatory sequences. Here, we employed our previously developed Gene Expression Engineering (GeneEE) platform for the creation of artificial 5^*′*^ regulatory sequences (ARES), containing both promoter and 5^*′*^ untranslated regions (UTR)^8^. GeneEE has been successfully utilized to generate functional ARES *in vivo* in various bacterial species including *Corynebacterium glutamicum, Escherichia coli, Pseudomonas putida, Streptomyces albus, S. lividans, Thermus thermophilus, Vibrio natriegens* and even Baker’s yeast^8,9^. This platform allows for the generation of highly diverse 200 nucleotide (nt) long random DNA sequences serving as promoter and 5^*′*^-UTRs, exceeding the total number and natural sequence variations observed in genomes for these regulatory elements.

The 200 nt long insert with the random DNA composition was cloned upstream of the RNA aptamer dBroccoli serving as a transcription reporter vector (see supplementary for plasmid maps). The RNA aptamer dBroccoli has the ability to bind tightly to a specific fluorogenic ligand ((Z)-4-(3,5-difluoro-4-hydroxybenzylidene)-1,2-dimethyl-1H-imidazol-5(4H)-one, DFHBI) that facilitates accurate and sensitive RNA detection, an essential feature in transcription analysis^10^. This aptamer is useful for the real-time observation of RNA synthesis and processing because its fluorescence upon ligand binding provides direct visualization of RNA molecules during transcription.

For efficient assembly of the plasmid library, we employed Golden Gate cloning^11^ (Figure 1A). To achieve a large library size, we performed multiple parallel transformations into chemically competent *E. coli* DH5α cells (Figure 1B). Following library construction, we confirmed the presence and correct size of the inserted DNA fragments using colony PCR with the GG-Col-F and GG-Col-R (for primer sequences please see Table S1) primer sets (Figure 1C) and Sanger sequencing (Figure 1D). Utilizing GeneEE, we constructed plasmid DNA libraries harboring approximately 1.54 million unique DNA templates.

**Figure 1.**
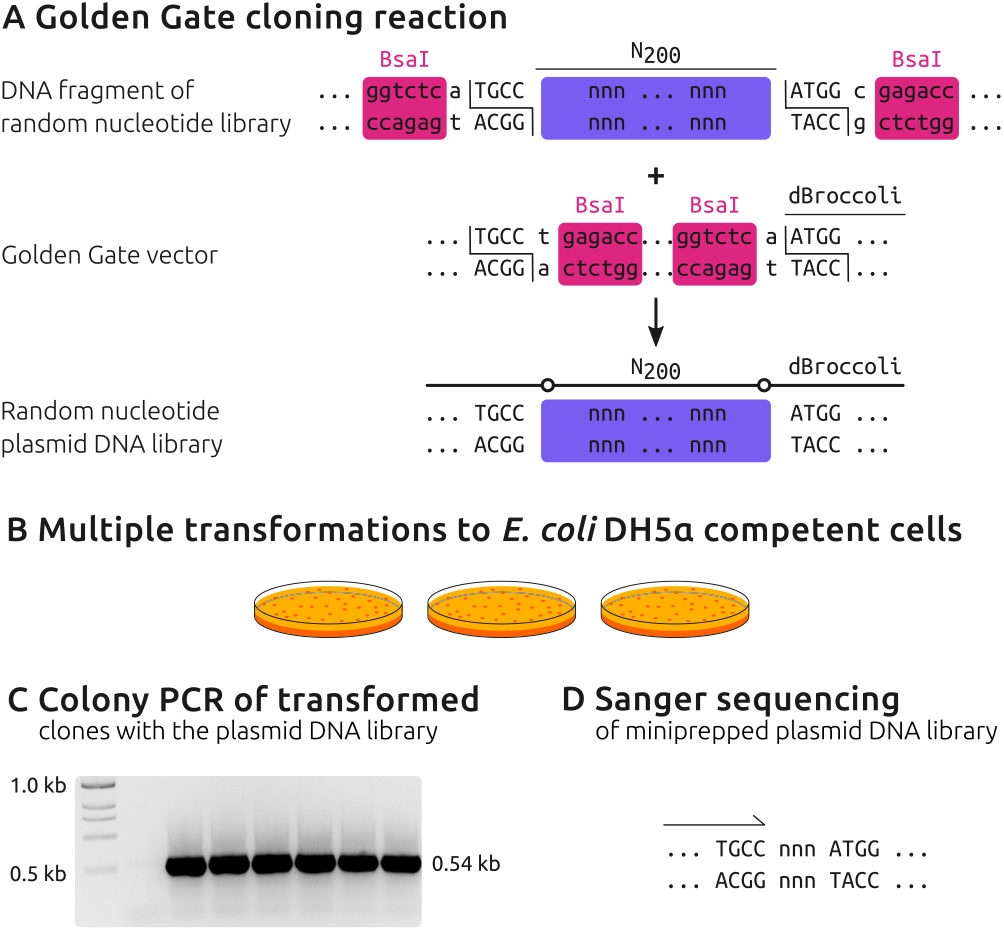
The schematic representation of the ARES plasmid DNA library construction. Golden Gate cloning using the Type II endonuclease, BsaI, to insert a 200 nt long ARES fragment into the backbone of a target vector (A). Multiple transformations of *E. coli* DH5α cells with a mixture of Golden Gate assembly reactions to achieve maximum diversity of ARES plasmid library (B). Agarose gel electrophoresis of the colony PCR-amplified product confirms the successful cloning of a 200 nt long ARES fragment in the plasmid backbone (C). Sanger DNA sequencing is further employed to confirm the successful cloning (D).

### DNA Sequencing

To identify the DNA template sequences within the extensive ARES plasmid DNA library, we employed NGS platforms from Illumina and Oxford Nanopore Technologies (ONT). Illumina MiSeq sequencing yielded a total of 15.8 million paired-end reads across two runs. Following quality filtering and dereplication using FLASH^12^, we retained 13.8 million high-quality reads for further analysis. ONT sequencing with GridION and PromethION flowcells provided an additional 11.4 million reads. After dereplication across all four libraries, a total of 6.4 million unique reads were obtained. Read abundance ranged from 1 to 109,425, with an average of 3.2 reads per sequence.

Clustering these unique sequences with VSEARCH^13^ at a 90% similarity threshold identified 1.54 million unique ARES sequences. Notably, 587,082 of these sequences were supported by at least three independent reads, indicating high confidence in their accuracy.

### *in vitro* Transcription Assays

The ARES plasmid DNA library served as templates in IVT reactions (Figure 2). These reactions utilized the core RNA polymerase and σ^54^ factor separately purified from *P. putida*, along with the activator ΔA2-DmpR following published protocols^14^. ΔA2-DmpR is a modified variant of the DmpR transcription regulator that lacks its regulatory A domain, rendering it independent of specific phenolic activation signal and thus constitutively capable of promoting transcription from the σ^54^-dependent *P*_O_ promoter it natively controls^15^. The DNA-binding domain of DmpR mediates high affinity to upstream activator sequences (UAS) located between –127 and –172 within the *P*_O_ promoter region. While the interaction between DmpR and core promoter elements is essential for initiating transcription, we used ΔA2-DmpR for its ability to promote transcription by σ^54^-RNAP in the absence of specific DNA binding sites^14^, which enabled unbiased analysis of σ^54^-dependent promoter activity across our library. A control experiment clearly demonstrated that RNA synthesis does not occur in the absence of ΔA2-DmpR, highlighting its crucial role in transcription initiation (Figure S1). After performing IVT-reactions, the resulting RNA was purified and sequenced for further analysis.

**Figure 2.**
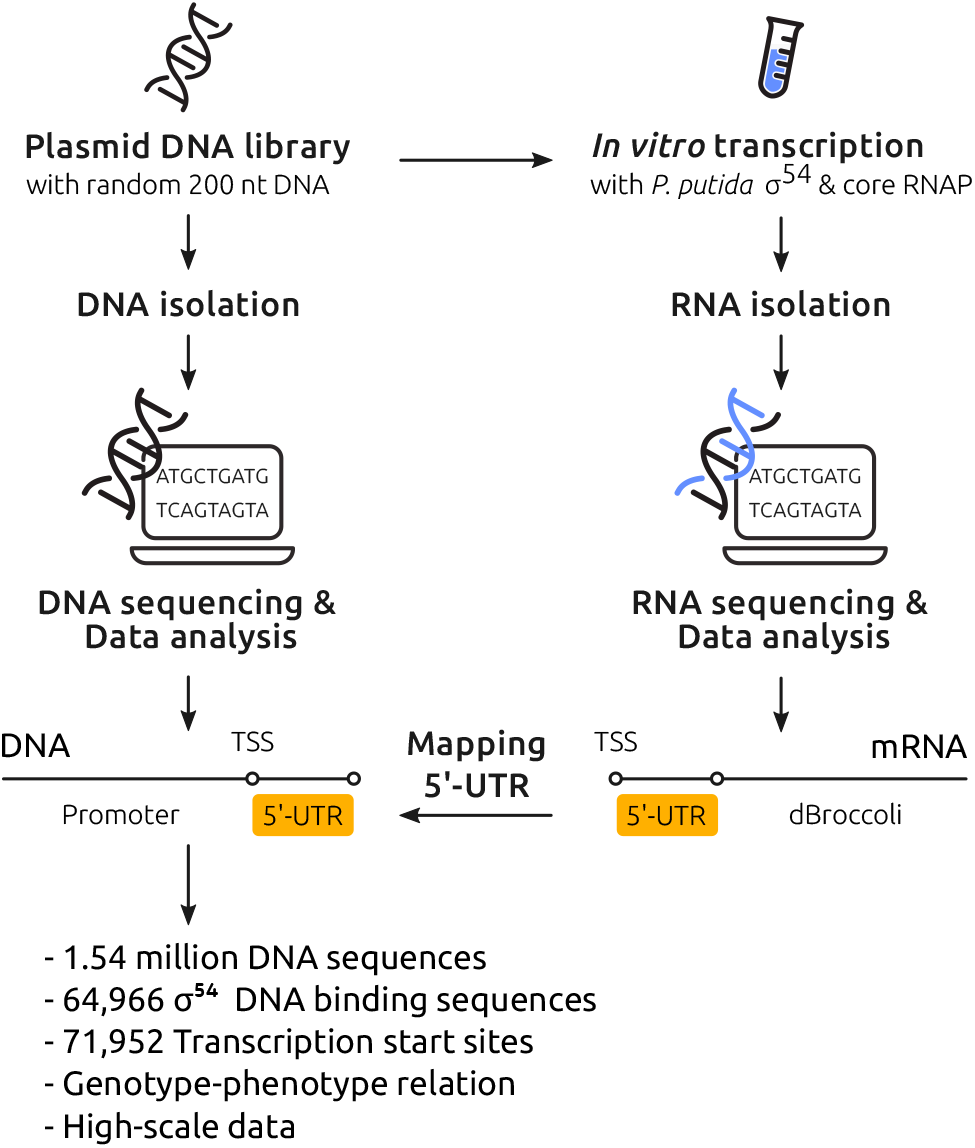
A schematic overview of the study. A random DNA plasmid library, containing ARESs to drive the expression of the RNA aptamer dBroccoli in relation to σ^54^-RNAP, was used during *in vitro* transcription assays. The generated RNA was isolated, sequenced, and mapped to the sequenced ARESs using 5^*′*^-UTR. *In silico* platforms were employed to identify *P. putida* σ^54^-associated promoter motifs. TSS, transcription start site

### RNA Sequencing Analysis

Deep sequencing of the 5^*′*^-enriched cDNA library generated from the *in vitro*-transcribed RNA yielded over 10 million reads. Following adapter trimming and alignment of these reads to the ARES sequences using BLASTn^16^, we identified over 2.7 million high-quality alignments corresponding to the template DNA. Filtering for reads mapped in the forward orientation and located within the defined ARES boundaries resulted in a set of 2.6 million valid alignments. These alignments identified a total of 152,992 potential transcription start sites (TSS). To enhance confidence, we applied a threshold of at least three reads mapping to a single position, resulting in a final set of 71,952 high-confidence TSS.

### Mapping RNA Transcripts to DNA Library

To identify the σ^54^ DNA binding sequences within ARES, we mapped the RNA transcripts back to their corresponding DNA templates. This process relied on the 5^*′*^-UTR sequence present in the transcribed RNA (Figure 2). By utilizing our knowledge of the DNA sequences of the RNA aptamer dBroccoli, the ARES—harbouring the 5^*′*^-UTR obtained from RNA sequencing—we were able to accurately link the 5^*′*^-UTR sequences in the RNA transcripts to their originating DNA templates within the ARES library.

Our constructed library contains approximately 1.54 million unique DNA templates. Although the cloning process initially used 200 nt single-stranded DNA sequences, DNA sequencing revealed that the ARES in the library range in size from 50 to approximately 250 nt, with the majority (around 0.8 million sequences) being 200 nt in length (Figure 3A). In total, 48,916 unique DNA templates were found to produce transcripts that could be mapped back to the ARES in the library.

**Figure 3.**
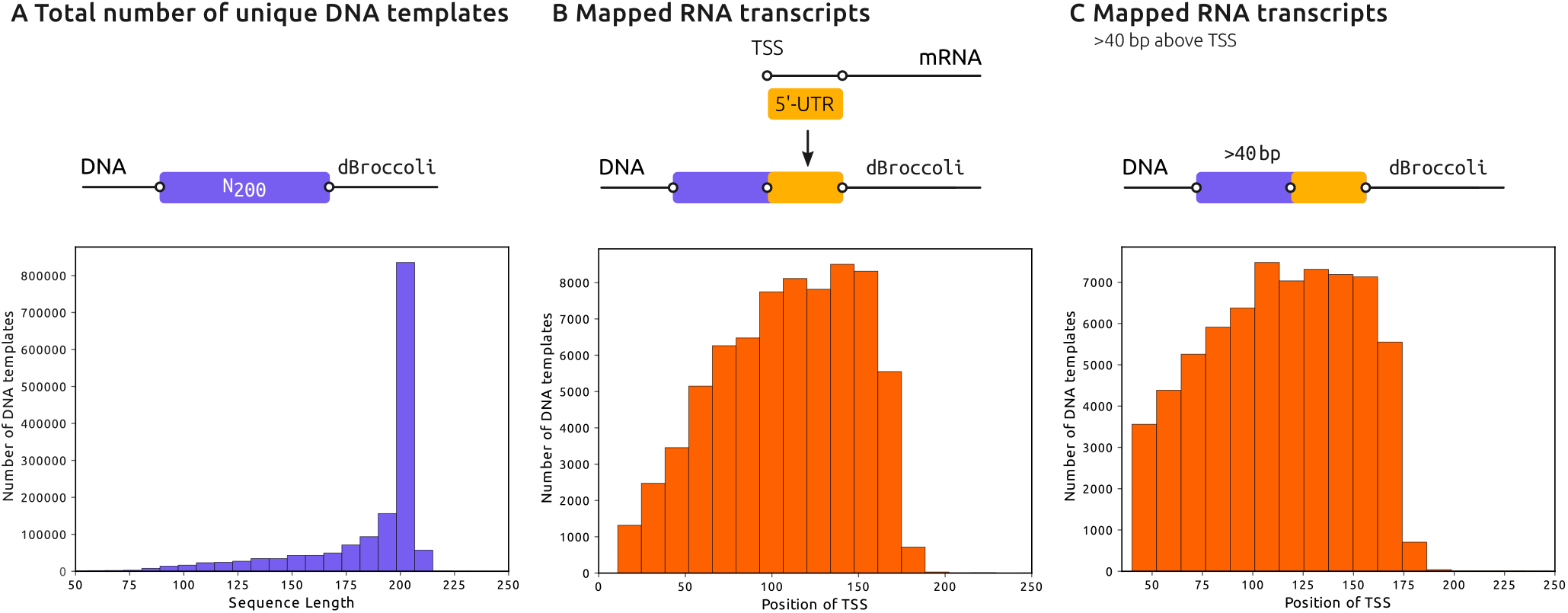
Overview of ARES Library and Transcript Mapping. A) Size distribution of ARES in the library. B) Distribution of TSS along the whole length of ARES. C) Distribution of TSS at or beyond position 40 within the ARES.

Some of the TSS were mapped to the very beginning of the ARES region, suggesting that certain sequences in the plasmid backbone may also function as weak promoters, allowing the σ^54^ transcriptional machinery to initiate transcription. The distribution of TSS along the entire length of the ARES is shown in Figure 3B. The σ^54^-RNAP recognizes promoters with conserved regions at positions –12 and –24 relative to the TSS^17^. Given their upstream location relative to the TSS, we established a threshold to include only those ARES where transcripts begin mapping at position 40 or beyond. After applying this threshold, 64,966 TSS were shortlisted. The distribution of these TSS, starting at or beyond position 40, is illustrated in Figure 3C.

### Identification of *P. putida* σ^54^-binding motifs

To identify the motifs in the 40 base pair (bp) region upstream of TSS in the ARES library, we used the Gibbs sampling algorithm BioProspector^18^. Our search focused on two motif regions at positions –12 and –24, each with a width of 6 bp, separated by a spacer ranging from 6 to 15 bp. These parameters were based on experimentally verified σ^54^ promoters in various species of *Pseudomonas*^19^.

The motif search was conducted on 48,916 unique DNA templates with 64,966 identified TSSs. This discrepancy arises because some ARES harbour multiple TSS. Several motifs were identified at both the –12 and –24 positions. While the exact positions of these two motifs are not strictly defined and their distance from the TSS vary, we still refer to them as the –12 and –24 positions, following the established naming convention. In the 40 bp stretch across all 64,966 templates, motifs were distributed as expected: the –12 motif was located from –26 to –5, peaking specifically at positions –16 to –14, whereas the –24 motif spanned from position –39 to –17, with a predominant presence from positions –29 to –26 (Figure 4A). For instance, in *E. coli* σ^54^ promoters, the average distance from the center of the –12 box to the TSS is 7 nucleotides, although this distance can vary from 1 to 12 nucleotides^20^. The spacer length between the two promoter boxes also varies, ranging from 15 to 21 nucleotides for a wide selection of σ^70^ endogenous promoters, but extending up to 23 nucleotides in synthetically designed promoters^21^.

**Figure 4.**
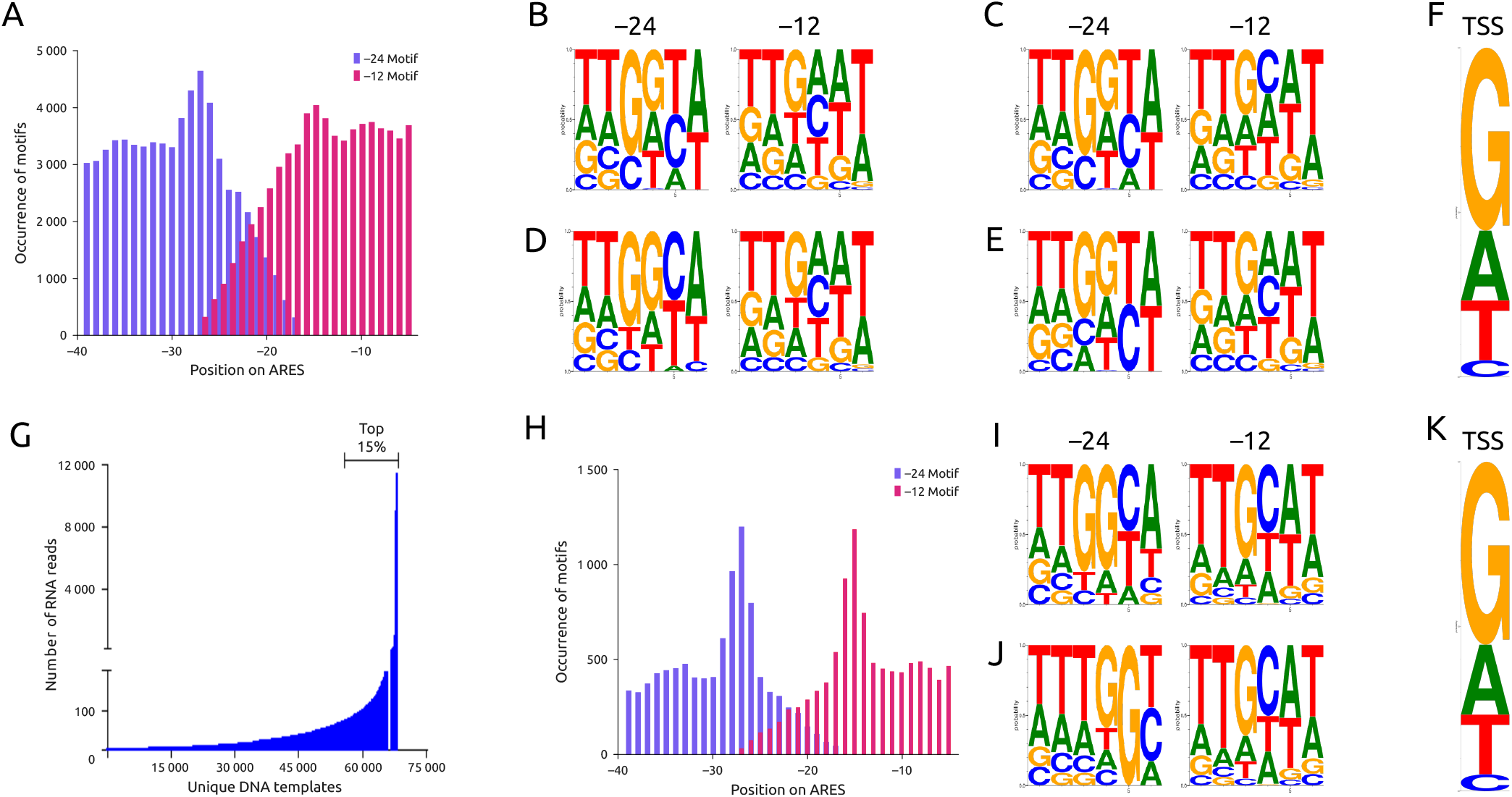
Comprehensive analysis of σ^54^ promoter motifs in *P. putida*. A) Distribution of motifs in a 40-bp stretch of ARES mapped with RNA. B-E) Sequence logos representing the refined σ^54^-specific –12 and –24 promoter motifs, derived from an extensive analysis of 64,966 unique DNA templates. F) Frequency analysis of nucleotide occurrences at the σ^54^ transcription start site (TSS) in *P. putida*. G) Graphical representation of the RNA read distribution across the ARES in the library. H) Distribution of motifs in a 40-bp stretch in the most highly transcribed ARES (top 15%) within the library. I and J) Two pair of -12 and -24 motifs identified in the high-expressing group. K) Frequency of nucleotides at their TSS.

BioProspector analysis identified combinations of both –12 and –24 consensus sequences. At the –12 position, two consensus sequences (TTGAAT and TTGCAT) were identified, while at –24, two consensus sequences (TTGGTA and TTGGCA) were observed (Figure 4B-E). In total, four pairs of these motifs were identified: TTGGTA/TTGAAT, TTGGTA/TTGCAT, TTGGCA/TTGAAT, and TTGGTA/TTGAAT (Figure 4B-E).

The RNA sequencing performed in this study also yielded quantitative data, including the count of RNA reads mapped to each ARES in the library. Figure 4G depicts how RNA reads are distributed across the shortlisted 64,966 unique DNA templates. The top 15% of ARES, comprising 9,451 sequences with high RNA read counts ranging from 45 to 11,469, were analysed using BioProspector to detect conserved motifs within the 40 bp region upstream of the TSS. Similarly, the motifs were predominantly located around the –12 and –24 positions (Figure 4H) for this dataset as well. However, unlike the broader search across all 64,966 templates, this targeted analysis on ARES leading to high levels of transcripts distinctly highlighted the two promoter boxes at their respective positions. At the –12 position, the consensus sequence (TTGCAT) was exclusively identified. At the –24 position, two consensus sequences (TTTGGT and TTGGCA) were noted. In conclusion, two motif pairs were distinguished: TTGCAT/TTGGCA and TTGCAT/TTTGGT (Figure 4I and J).

In addition to the presence of core promoter motifs, UASs are associated with binding to the transcriptional activator DmpR^15^. In this study, we employed ΔA2-DmpR, which can activate transcription without requiring specific UAS interaction^14,22^. Although ΔA2-DmpR does not bind to its specific UAS within the current experiment, we also sought to explore unique sequence patterns in regions beyond the previously analyzed 40 bp segment for core promoter motifs. We performed a motif search for the occurrence of a single conserved motif within a 97 bp region upstream of the TSS in the ARES library, for motifs ranging from 6 to 11 nucleotides in length. For the top 15% of ARES, composed of 5,767 sequences (the increase in the upstream sequence length from 40 to 97 nucleotides reduced the number of templates from 9,451 to 5,767), known for high transcript expression *in vitro*, our search for a 6 bp motif identified the consensus sequence (TTGCAT) previously noted at the –12 position in core promoter analysis, but peaking at the –17 position (Figure S2a). By incrementally extending these motifs by one nucleotide, the motifs consistently appeared at the same location, displaying almost identical sequences with an added T nucleotide at the beginning (Figure S2b-f). Furthermore, an analysis within the 40 bp region upstream of the TSS among the highest expressing ARES subset reaffirmed the presence of the same motif (TTGCAT), aligning with earlier findings from core promoter studies (Figure S3).

In this study, we also conducted an extensive frequency analysis to assess nucleotide occurrences at the σ^54^ TSS in *P. putida*. Our analysis indicates that guanine (G) is the most frequent nucleotide, found in approximately 55% of the ARES within our library, as shown in (Figure 4F and K). This prevalence of G at the TSS is consistent with findings from previous studies on *Pseudomonas* species. Genes such as *algC, algD, cpg2, flesR, oprE*, and *xylS*, which are all regulated by σ^54^, typically show a high occurrence of G at the TSS. Further comparative analyses of σ^54^-dependent promoter sequences across various bacterial species also consistently identify G as the dominant nucleotide at the TSS^17^.

### IVT-based validation of transcript-producing ARES

To validate the functionality of positive ARES that showed an affinity for binding with the σ^54^-RNAP complex and generating transcripts, we performed IVT reactions with reconstructed plasmid DNA. For this validation experiment, 18 ARES, each 200 nt long (Table S2), were randomly selected and cloned into the Mango-pSEVA2311 plasmid. The transition from the dBroccoli to the RNA aptamer Mango III was motivated by two main factors: Firstly, the Mango III aptamer offers a superior signal-to-noise ratio^23^, enhancing the quality of readouts across a wide range of promoter activities from low to high. Secondly, it is more suitable for microfluidics-based IVT reactions, which are concurrently being conducted within the same project. We conducted parallel experiments using both negative and positive controls. The negative control consisted of all reaction components except the DNA template, while the positive control used the σ^54^-dependent *P*_O_ promoter^22^, which demonstrated the highest transcriptional activity, as measured by the fluorescence intensity of the RNA aptamer Mango III (Figure 5A and B).

**Figure 5.**
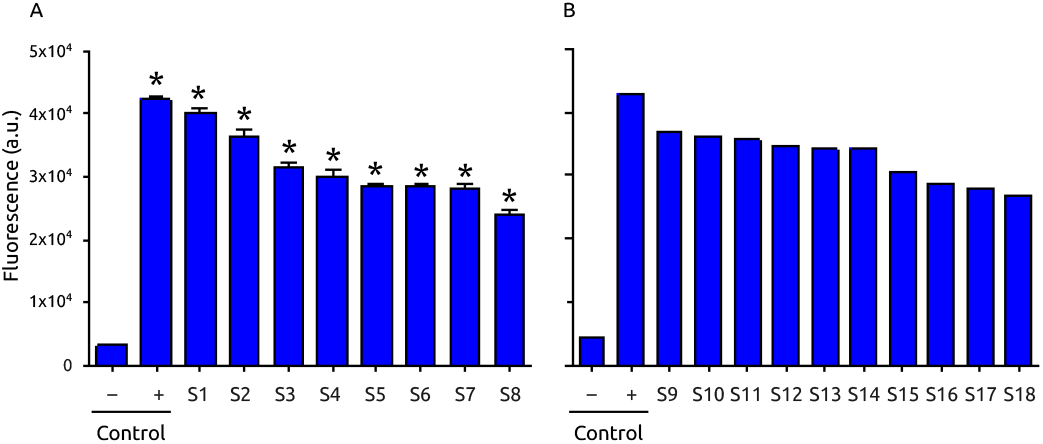
Evaluation of transcript-producing ARES in IVT reactions. A) Fluorescence intensity measurements of the RNA aptamer Mango III for the first eight randomly selected ARES, performed in triplicate alongside both positive and negative controls. Data are presented as mean ± s.d. for n=3 technical replicates. Bars marked with asterisks (*) indicate statistically significant differences from the Negative Control. B) Fluorescence intensity of the RNA aptamer Mango III for the remaining ten randomly selected ARES, each measured in a single reaction.

To evaluate the statistical significance of transcript production, we conducted a statistical analysis. An ANOVA test produced an F-statistic of 738.84 (*p* < 0.0001), indicating a strong variance among the groups. Post-hoc comparisons using Tukey’s honestly significant difference test showed that all experimental conditions, including the Positive Control and various test groups (S1, S2, S3, S4, S5, S6, S7, S8), differed significantly from the Negative Control, with adjusted *p*-values below 0.001 for all comparisons. These findings are visually presented in Figure 5A, where mean values, and standard deviations are marked with asterisks to indicate statistically significant differences from the Negative Control.

### Predicting the occurrence of IVT generated σ^54^-RNA polymerase-binding motifs in the *Pseudomonas putida* KT2440 genome

Building on the insights gained so far, we sought to explore how these findings can be applied to annotate the genome of *P. putida* KT2440. Our search revealed that the only online database currently annotating σ^54^ promoters is the BioCyc genome database collection^24^. However, within this database, the annotation for *P. putida* KT2440 includes only ten σ^54^ promoters (as seen in BioCyc_rpoN). This limited annotation underscores the need for a more comprehensive approach to identifying σ^54^-RNA polymerase-binding motifs across the genome. For this purpose, we used the Find Individual Motif Occurrences (FIMO) software^25^. We applied FIMO to search a 150 nt stretch upstream of all coding sequences (CDSs) in the *P. putida* KT2440 genome (Genome accession number: NC_002947.4) for σ^54^-RNA polymerase-binding motifs reported in this study. This software calculates a log-likelihood ratio score for every position in a specified sequence dataset, utilizing dynamic programming techniques to convert this score into a *p*-value^26^. A stringent *p*-value threshold of 0.005 was set to determine the significant presence of σ^54^ DNA-binding motifs.

Using this criteria and considering a spacing of four to seven nucleotides between the –12 and –24 motifs, 68 genes were identified (Table S3). A sub-selection of the identified genes, along with their molecular functions and the sequences of the –12 and –24 motifs located within the 150 nt stretches, is displayed in Table 1. While σ^54^-regulated genes in *P. putida* are not extensively documented^27,28^, several of the genes we identified have been reported to be regulated by σ^54^ in other bacterial species, such as *Bacillus subtilis, Desulfovibrio vulgaris, Escherichia coli, Paraburkholderia phumatum, Pseudomonas aeruginosa, Vibrio parahaemolyticus*, and *Xanthomonas oryzae*^29–35^.

**Table 1.** Significant annotation of IVT-generated σ^54^-RNA polymerase binding motifs in the upstream regions of genes involved in various biological processes.

| Gene | Product | –24 Motifs | –12 Motifs | References |
| --- | --- | --- | --- | --- |
| **Flagellar assembly** |  |  |  |  |
| *flgA* | Flagellar basal body P-ring formation chaperone FlgA | TCGGCA | TTGCTT | 27,34 |
| *flgB* | Flagellar basal body rod protein FlgB | TTGGCA | TTGCTA | 27,34 |
| *flgF* | Flagellar basal-body rod protein FlgF | TTGGTT | TTGCTT | 27,34 |
| *flgG* | Flagellar basal-body rod protein FlgG | ATGGCT | TTGCAA | 34 |
| *fliE* | Flagellar hook-basal body complex protein FliE | CTGGCA | TTGCTT | 27 |
| *motA* | Flagellar motor stator protein MotA | TTGGTT | TGGCAT | 28 |
| **Metabolism** |  |  |  |  |
| PP_2783 | 3-oxoacyl-ACP reductase family protein | TGGGCT | TGGCAT | 35 |
| *eutC* | Ethanolamine ammonia-lyase subunit EutC | TTGGCA | TTGCAT | 33 |
| PP_1786 | Glycosyltransferase | ATGGCA | TTGCAT | 32 |
| PP_0859 | Amidohydrolase | CTGGTA | GTGCTT | 30 |
| *edd* | Phosphogluconate dehydratase | GTGGCA | TTGTTT | 34 |
| PP_2836 | Fumarylacetoacetate hydrolase family protein | GTGGCT | TTGCAA | 27 |
| PP_1946 | Glucose 1-dehydrogenase | TTGGCA | TTGCAA | 27 |
| PP_3486 | Cytochrome c | ATGGCT | GTGCTT | 34 |
| PP_1141 | ABC transporter substrate-binding protein | TGGGCT | TGGCAT | 34 |
| *ppx* | Exopolyphosphatase | ATGGCA | TTGCAT | 36 |
| PP_3778 | Pyrroline-5-carboxylate reductase | TTGACA | TTGCTA | 29 |
| *xdhA* | Xanthine dehydrogenase small subunit | TGGGCA | GTGCAT | 31 |
| **Transport** |  |  |  |  |
| PP_4970 | Cytochrome c | GTGGCT | GTGTTT | 34 |
| *ntrB* | Nitrogen regulation protein NR(II) | GTGGTT | TTGCAT | 34 |
| **Response to stress** |  |  |  |  |
| PP_3248 | Dyp-type peroxidase | TTGATA | TTGCAA | 32 |
| PP_3539 | MerR family DNA-binding transcriptional regulator | GTGGTT | TTGAGT | 34 |
| **Transcription regulation** |  |  |  |  |
| PP_2952 | LysR family transcriptional regulator | GTGGTT | TGGCAT | 34 |
| PP_0952 | RNA polymerase factor sigma-54 | AAGGCA | TTGCTT | 34 |
| **Uncharacterized** |  |  |  |  |
| PP_1450 | HlpA activation/secretion protein HlpB | GTGGCT | TAGCAT | 34 |

The σ^54^ complex is known to regulate the expression of flagellar genes to facilitate bacterial motility^34^. Our findings confirm that the σ^54^-RNA polymerase binding motifs identified via ARES-mediated IVT prominently locate in the promoter regions of flagellar genes in *P. putida* KT2440 (Table 1). Furthermore, genes encoding metabolic proteins involved in various processes like nucleic acid metabolism, glucose metabolism, oxidative phosphorylation, and amino acid metabolism are regulated by the σ^54^-RNA polymerase complex, aligning with the observations in other bacterial species^27,29–32,36^. Promoters of genes linked to transport, transcription regulation, and stress response in *P. putida* KT2440 also contain significant σ^54^-RNA polymerase binding motifs (Table 1). The method of using FIMO software to map ARES-generated motifs sets a new standard in the predictive identification of σ-factor binding motifs within bacterial genomes, enhancing our understanding of the complex roles σ-factors play in regulating bacterial genes and metabolisms.

## Discussion

Bacterial σ-factors direct RNA polymerase to specific promoter regions, thereby directly influencing gene expression and cellular responses. The evolutionary adaptation of these nucleotide preferences demonstrates how bacteria respond to environmental pressures, resulting in a diversification of regulatory sequences and strategies across different species. These insights into the promoter specificity of σ-factors are invaluable, not only for understanding transcriptional regulation in bacteria but also for advancing biotechnological applications. This includes the design of synthetic promoters to optimize gene expression and the development of promoter prediction tools.

Our study leveraged an extensive DNA library, encompassing over 1.54 million DNA templates, to explore σ-factor interactions across a wide array of constructs. A key outcome of this study sets the groundwork for expanding the research to include other σ-factors from diverse bacterial species enabling the generation of high-quality data that is suitable for training machine learning models for developing among others transcriptional models, promoter prediction tools and re-annotation of genomes. By including both functional and non-functional transcription sequences—the latter often overlooked in datasets—we enhance the training of computational models to discern functional from non-functional DNA sequences. Such an inclusive approach is crucial for developing more precise predictive models in computational biology, offering a deeper and more holistic understanding of the elements that govern transcriptional regulation. The versatility of this method promises to generate further high-quality data, broadening our comprehension of σ-factor dynamics across various bacterial contexts.

As the field of genomics advances, there is a growing need to expand the capabilities of automated genome annotation tools beyond the prediction of CDSs alone. Current automated genome annotation tools predominantly focus on identifying CDSs, leaving regulatory sequences underexplored. Developing additional tools that can accurately predict and annotate these regulatory sequences is, therefore, crucial for a comprehensive understanding of genome functionality.

In this realm, our approach offers a robust solution for predicting regulatory sequences, significantly enhancing genome annotations. This integration provides a more nuanced understanding of the regulatory landscapes of bacterial genomes, paving the way for future research to explore uncharacterized regions of the genome and potentially uncover novel aspects of bacterial life. By identifying and annotating regulatory elements, we can extend our comprehension of genome functionality, opening new avenues for genetic and microbial research.

## Materials and Methods

### Strains and growth conditions

*E. coli* DH5α was cultivated in lysogeny broth medium containing 50 mg/L kanamycin for library plasmids (pHH100-dBroccoli vector).

### Composition of the single-stranded random nucleotide oligo

To create the extensive promoter DNA libraries, a 200 nt long stretch of single-stranded Random Nucleotide Oligomers (ss-RaNuqO) was procured from Integrated DNA Technologies (IDT), Inc. (Belgium) in the form of a four nanomole ultramer. The ss-RaNuqOs are equipped with adapters on both ends. Adapter1 carries the BioBrick Prefix, while Adapter2 contains the BioBrick Suffix sequences. Each adapter also incorporates a type IIS restriction enzyme recognition sequence for BsaI (5^*′*^-GGTCTC(N1)/(N5)-3^*′*^), accompanied by overhang sequences TGCC in Adapter1 and TACC in Adapter2. These adapters also provide the complementary DNA strand for primers to make double-stranded random nucleotide DNA through polymerase chain reaction (PCR). For further details please see Lale, et al^8^.

### Generation and cloning of double-stranded 200 nt long 5^*′*^ artificial regulatory sequences

To create the double-stranded random nucleotide insert, a 12-cycle PCR was employed, utilizing the ss-RaNuqO as the DNA template and the BBa-Prefix-F and BBa-Suffix-R as primers (Table S1). The selection of a limited number of cycles was deliberate, to avoid the potential PCR amplification bias that could otherwise result in reduced sequence diversity.

The double-stranded 200 nt long insert was cloned upstream of the dBroccoli aptamer in the pHH100 vector using Golden Gate cloning. The dBroccoli aptamer was initially extracted from the pTRA51hd plasmid using the BB1-F and BB1-R primers, which include overhangs for Gibson assembly into the pHH100 backbone. The amplified aptamer was subsequently cloned into the pHH100 backbone, which had been pre-amplified with the BB2-F and BB2-R primers. This cloning process was carried out using the Gibson assembly method with the NEBuilder HiFi DNA Assembly Master Mix. The pHH100-dBroccoli backbone was initially amplified with the BB3-F and BB3-R primer sets, which incorporated a type IIS restriction enzyme recognition sequence for BsaI. This sequence was accompanied by overhang sequences ACGG and ATGG, which are complementary to TGCC in Adapter1 and TACC in Adapter2 of the artificial promoters, respectively. The plasmid DNA library was transformed into *E. coli* through a heat shock procedure. The library consisted of 10^6^ clones. Colonies were scooped from LB agar plates and ARES plasmid library was isolated following the QIAprep Spin Miniprep Kit (Qiagen) protocol.

### Phenol-Chloroform purification of the plasmid DNA library

The Miniprep DNA was purified using the Phenol-Chloroform purification protocol. To begin, a one-volume mixture of phenol, chloroform, and isoamyl alcohol (25:24:1) was added to the sample. This mixture was vortexed for 20 seconds and then centrifuged at room temperature for 5 minutes. The upper aqueous phase was extracted and transferred to a new tube, ensuring no phenol contamination. The DNA in the aqueous phase was precipitated at –20 °C overnight by adding glycogen (20 μg), ammonium acetate (3.75 M), and 100% ethanol (2.5 times the combined volume of both the sample and ammonium acetate) in the specified order. After precipitation, the sample was centrifuged to pellet the DNA, the supernatant was removed, and two ethanol (70%) washes were performed. Ethanol was thoroughly removed, and the purified DNA was resuspended in nuclease-free water for *in vitro* transcription assays.

### *In vitro* transcription with *P. putida* σ^54^-RNA polymerase

The *in vitro* transcription assays were conducted at 30 °C in an acetate buffer (AcB) with a composition of 35 mM Tris-acetate (pH 7.9), 70 mM potassium acetate, 20 mM ammonium acetate, 5 mM magnesium acetate, 1 mM DTT, and 0.275 mg/ mL bovine serum albumin. These assays employed 3 mM ATP and 15 nM supercoiled ARES library plasmid, following established protocols. The final reaction mixture also included *P. putida* core RNA polymerase (80 nM), σ^54^ (400 nM) and ΔA2-DmpR (400 nM). Prior to the addition of ATP and the DNA template, the core RNA polymerase and σ^54^ were pre-incubated for 5 min at 30°C to facilitate holoenzyme formation. Subsequently, the reaction was incubated with the DNA template and ΔA2-DmpR for an additional 20 min to allow closed-complex formation. Transcription was initiated by introducing NTPs, with final concentrations of 360 nM each for GTP, CTP, ATP, and UTP. Heparin (0.1 mg/mL) was used to prevent re-initiation. To terminate the reaction, 5 μL of a stop load mix (comprising 150 mM EDTA, 1 M NaCl, 14 M urea, 3% glycerol, 0.075% (w/v) xylene cyanol, and 0.075% (w/v) bromophenol blue) was added. Finally, dBroccoli fluorescence was quantified using a Tecan Infinite M1000 Pro Plate Reader.

An IVT assay was also conducted to validate the ARES of a library that generated transcripts containing a promoter sequence for *P. putida* σ^54^. Following the sequencing analysis of both the ARES DNA template and IVT-generated transcripts, eighteen candidates were selected, named p1 to p18 (Table S2). The complete 200 nt sequence was cloned upstream of the Mango III aptamer within the pSEVA2311 plasmid, utilizing Golden Gate cloning. Comprehensive details on the primers used for this cloning process, designated as p1 to p18 for both the fragment (Frag) and the backbone (BB), can be found in the supplementary section, Table S1. Before this, the Mango III aptamer, synthesized by Twist Bioscience, was amplified using the Lib1-F and Lib2-R primers. This amplified aptamer was then cloned into the pSEVA2311 backbone, which had been previously amplified with the Lib3-F and Lib4-R primers, through the Gibson assembly process, employing the NEBuilder HiFi DNA Assembly Master Mix. Successful assembly of the Mango-pSEVA2311 constructs was confirmed through colony PCR with the Lib7-F and Lib8-R primers. To ensure high-quality templates for the IVT reaction, the resulting constructs were purified using a phenol-chloroform extraction method, and subsequently used for the IVT reaction as previously described.

### Purification of *in vitro* transcribed RNA

Plasmid DNA was eliminated from the RNA samples using the Turbo DNAse Kit (Invitrogen). Two successive 15 minute DNAse (2 units) treatments were conducted immediately after transcription. Following the second treatment, RNA purification was carried out according to Monarch RNA Cleanup Kit protocol (New England Biolab), and quantification was performed using Nanodrop (ThermoFisher).

### Sequencing of ARES plasmid library

DNA sequencing was done by creating a custom sequencing library by first amplifying the 200N stretch with the SeqDNA forward primer and the SeqDNA reverse primer using Phusion DNA polymerase (New England Biolabs) for 15 cycles, followed by 10 cycles PCR using again Phusion DNA polymerase and TruSeq index primers. Sequencing was performed using two different sequencing technologies. On the one hand, a MiSeq sequencing platform (Illumina) with a 600 cycle MiSeq Reagent Kit v3 (Illumina) was used to generate 2x 300 nt paired end reads. The Illumina reads were trimmed using CUTADAPT^37^ to trim the 3^*′*^-ends with option -a ATGGACGAGC for the R1 reads and -a GGCATCCGAC for the R2 reads, followed by joining of the forward and reverse reads using FLASH^12^ with parameters -m 50 -M 295. After read joining, the 200N stretch as well as 10 nt up- and downstream were extracted using CUTADAPT run with parameters –discard-untrimmed, -a GGCATCCGAC…ATGGACGAGC, and –action retain. On the other hand, the GridION sequencing platform (Oxford Nanopore Technologies) was used to sequence a LSK109 library on an R9.4.1 flowcell and the PromethION platform to run a LSK112 library on an R9.4.1 flowcell. Afterwards, the read data was filtered using FASTP^38^ to remove reads shorter than 250 nt, longer than 600 nt, or with a quality below 5. The remaining reads were trimmed using two chained CUTADAPT runs with parameters –discard-untrimmed, -g GGCATCCGAC, and –action retain respectively –discard-untrimmed, -a ATGGACGAGC, and –action retain to extract the 200N stretch as well as 10 nt up- and downstream.

All data sets were then de-replicated individually using VSEARCH^13^ in mode --derep_fulllength with the parameters --sizeout. Afterwards, the resulting files were combined, dereplicated again (parameters: --sizeout --sizein), and finally clustered and merged into ARES consensus sequences via --cluster_size with parameters --sizeout --sizein --id 0.90 --clusterout_sort --minseqlength 120. The consensus sequences were finally renamed consecutively based on read abundance.

All ARES consensus sequences without the leading and trailing 10 nt of the vector are available via BioProject PRJNA1054479, SRA accession SRR27486701.

### Sequencing of *in vitro* transcribed RNA

For RNA sequencing, the following protocol was performed: First, the RNA was capped using the RNA capping reagents and protocol M2080 from New England Biolabs. Second, reverse transcription and template switching with resulting capped RNA was done using two biotinylated oligos (CCGACCGTCTCAGATGGACC and ACTCGACATCTGAGCCCAC) with the Template Switching RT Enzyme Mix M0466 (New England Biolabs) and the DNA-RNA TS-oligomer CCCTACACGACGCTCTTCCGATCGArGrG+G (IDT). Third, after first strand synthesis, the RNA-DNA hybrid was first purified using the QIAGEN MinElute kit according to the manufacturer’s instructions with an elution volume of 25 μL, followed by coupling to 25 μL M280 streptavidin-coated magnetic beads (Invitrogen) prepared according to the manufacturer’s protocol. The beads were then washed twice with 100 μL 0.1M NaOH (1 min incubation), twice with 200 μL 1x binding and wash buffer, twice with 200 μL water and finally resuspended in 25 μL water. Fourth 10 μL bead suspension were used for cDNA amplification using the SeqRNA forward primer and the SeqRNA1 reverse primer and SeqRNA2 reverse primer. Finally, the PCR product was used in conjunction with the SQK-LSK114 library kit (Oxford Nanopore Technologies) to prepare a sequencing library that was run on a PromethION R10.4.1 flowcell.

### Mapping of transcribed RNA with ARES plasmid library

The reads were preprocessed using CUTADAPT with parameters –discard-untrimmed and -g CCCTACACGACGCTCTTC-CGATCGAGGG to remove the TSO sequence. For mapping to the ARES consensus sequences MINIMAP2 was used with parameters -a -x map-ont --secondary=no --sam-hit-only. The results were filtered to remove hits not mapping in forward direction and those mapping starting in the vector sequence. Together with the FASTA file containing the ARES sequences, the filtered read data were then used as input for a custom Perl script (MARKTSS.PL) to identify each position with at least 3 mapped reads.

### Data handling and analysis of promoter motifs in the ARES of the library

Data analysis was conducted using Python, leveraging various libraries such as BioPython, Pandas and NumPy for efficient processing, manipulation, and bioinformatics-specific tasks. The function fasta_to_dataframe of biopython library serves as a useful tool for converting bioinformatics data in FASTA format into a structured tabular form for further analysis and manipulation using Pandas. This function extracted metadata from sequence identifiers, including centroids, sequences, sizes, positions, and mapping statuses, and compiled them into a Pandas DataFrame. The to_parquet function of Pandas, enabling the storage of DataFrames in the efficient Parquet format. This columnar storage file format is specifically designed for optimal performance within big data processing frameworks. Another function, pd.read_parquet excels in efficiently reading data stored in the Parquet file format.

BioProspector, a C program, was utilized to analyze the upstream region of TSS in the ARES of a library, searching for promoter sequence motifs. BioProspector employs background models, ranging from zero to third-order Markov models, with parameters that are customizable by the user or can be estimated from a specified sequence file^18^. In the present study, the parameters were taken from experimentally verified σ^54^ promoters in various species of *Pseudomonas*^19^. The significance of identified motifs is evaluated through a motif score distribution, determined using a Monte Carlo method^18^.

The parsing of motifs involved identifying blocks and patterns within the sequence data, achieved through a custom parsing function. This function interpreted lines from input files to extract relevant information about positions, blocks, and consensus sequences. The parsed motifs were then converted into comprehensive DataFrames, which provided detailed overviews of the motifs, including their positions, consensus sequences, and degeneracies. Sequence logos were generated using the seqlogo library to visualize motif patterns, while additional visualizations were created using Matplotlib to represent data distributions and relationships effectively.

## Data availability

All the data and scripts are deposited in the git repository (https://github.com/LaleLab/Publication_Sigma54).

## Acknowledgements

We acknowledge the funding from the Research Council of Norway (grant no. 316129); NTNU-Biotechnology, Enabling Technologies Program (personal PhD stipend to L.T.); Faculty of Natural Sciences at NTNU (personal PhD stipend to M.F-L.). We thank Katharina Pflüger-Grau, Ph.D. for providing the plasmid pTRA-51hd harbouring the dBroccoli.

## Contributions

E.A.K. & R.L. conceived the study. E.A.K., C.R-R., G.S.D., L.T., M.F-L., T.B., J.K., V.S., & R.L. carried out data collection and analyses. E.A.K., C.R-R., & R.L. draw the figures. All authors wrote and corrected the paper.

