## Supporting Online Material for "High-resolution mapping of Sigma Factor DNA Binding Sequences using Artificial Promoters, RNA Aptamers and Deep Sequencing"

Khan EA et al, 2024.

**Table S1:** Primers used in this study

| Primer name | Sequence |
|---|---|
| **Cloning of double-stranded 200 nt long ARES to pHH100-dBroccoli backbone** | |
| BBa-Prefix-F | GAATTCGCGGCCGCTTCTAGAG |
| BBa-Suffix-R | CTGCAGCGGCCGCTACTAGTA |
| BB1-F | TCGCACGATATACAGGATTTTGCGGATTTGTCCTACTCAGGAG |
| BB1-R | GTCAGCTTTATGCTTGTAAACCGACCGAACAGGCTTATGTC |
| BB2-F | CGGTTTACAAGCATAAAGCTG |
| BB2-R | GCAAAATCCTGTATATCGTGC |
| BB3-F | CGTAATCAAGCCACTTCCTTTT |
| BB3-R | GTACGCGTACGGTCTCTGGCATCCGACACCCTGCGTCAATG |
| GG-Col-F | GCGGATTTGTCCTACTCA |
| GG-Col-R | CTGGATCATCAGAGTATGTG |
| **For validation, 18 ARES, each 200 nt long cloned into the Mango-pSEVA2311 plasmid** | |
| p1-Frag-F | GGCTACGGTCTCCAAGATATCCCGTATAACATTTGTACACTCCGTTATGG |
| p1-Frag-R | GGCTACGGTCTCTTGGCAACATTTACATTCAGCAGGCACCAGCGCA |
| p1-BB-F | GGCTACGGTCTCTGCCATGTGTATGTGGGCG |
| p1-BB-R | GGCTACGGTCTCCTCTTAATTAAAGGCATCAAATAAAACGAAAGG |
| p2-Frag-F | GGCTACGGTCTCGACGACTGTCTTAAGGTTAAG |
| p2-Frag-R | GGCTACGGTCTCACCACGAGACATATACAATTAC |
| p2-BB-F | GGCTACGGTCTCAGTGGATGTAAATGTTGCCATGTG |
| p2-BB-R | GGCTACGGTCTCGTCGTTAATTAAAGGCATCAAATAAAACG |
| p3-Frag-F | GGCTACGGTCTCCTTCGTGATAATGAGCTGAG |
| p3-Frag-R | GGCTACGGTCTCGGGGTAAAATTATATTTAAAGCCG |
| p3-BB-F | GGCTACGGTCTCGACCCCTGATGTAAATGTTGCCATGTG |
| p3-BB-R | GGCTACGGTCTCCCGAAATTTAATTAAAGGCATCAAATAAAACG |
| p4-Frag-F | GGCTACGGTCTCGTAATTGGATTTATTCTACTCTTCC |
| p4-Frag-R | GGCTACGGTCTCTCGAAAGAATATCACAATGTAG |
| p4-BB-F | GGCTACGGTCTCTTTCGGGGGTATGTAAATGTTGCCATGTG |
| p4-BB-R | GGCTACGGTCTCGATTATAACATTAATTAAAGGCATCAAATAAAACG |
| p5-Frag-F | GGCTACGGTCTCGCCTTTAATTAAGTTGTCCTGATTATGGTAGTACTAGTAG |
| p5-Frag-R | GGCTACGGTCTCTTGGCAACATTTACATATGCCCCGCCTCATCATATC |
| p5-BB-F | GGCTACGGTCTCTGCCATGTGTATGTGGGCG |
| p5-BB-R | GGCTACGGTCTCGAAGGCATCAAATAAAACGAAAGGC |
| p6-Frag-F | GGCTACGGTCTCTGAGATTCATTAAATATTATTGGGGAAGAATTGG |
| p6-Frag-R | GGCTACGGTCTCATGGACACCACCGTATCCGATG |
| p6-BB-F | GGCTACGGTCTCATCCATGTAAATGTTGCCATGTG |
| p6-BB-R | GGCTACGGTCTCTTCTCACTTTTAATTAAAGGCATCAAATAAAACG |
| p7-Frag-F | GGCTACGGTCTCACCTTTAATTAATAAGCGAGATTCAAGAATTTTTAG |
| p7-Frag-R | GGCTACGGTCTCCTAGTATAATTCCATTCCACAG |
| p7-BB-F | GGCTACGGTCTCCACTACACGCTGATGTAAATGTTGCCATGTG |
| p7-BB-R | GGCTACGGTCTCAAAGGCATCAAATAAAACGAAAG |
| p8-Frag-F | GGCTACGGTCTCCAAGGATTTTGAATAAAGTGGTAAAG |
| p8-Frag-R | GGCTACGGTCTCACAACATTTACATCCGCTATATGGCTAAATTAG |
| p8-BB-F | GGCTACGGTCTCAGTTGCCATGTGTATGTGG |
| p8-BB-R | GGCTACGGTCTCCCCTTTAATTAAAGGCATCAAATAAAACG |
| p9-Frag-F | GGCTACGGTCTCTGAGTTTACTTCGGGTAAC |
| p9-Frag-R | GGCTACGGTCTCCCACATGGCAACATTTACATCTACCTATACCACCTAAGAG |

| | |
|---|---|
| p9-BB-F | GGCTACGGTCTCCTGTGTATGTGGGCGTACG |
| p9-BB-R | GGCTACGGTCTCTACTCCATTTAATTAAAGGCATCAAATAAAACGAAAG |
| p10-Frag-F | GGCTACGGTCTCAAAAGAGTATGATCCCATTGG |
| p10-Frag-R | GGCTACGGTCTCGACCATTGATAGGATAGAAAATAAC |
| p10-BB-F | GGCTACGGTCTCGTGGTTTGAATGTAAATGTTGCCATGTG |
| p10-BB-R | GGCTACGGTCTCACTTTAATTAAAGGCATCAAATAAAACG |
| p11-Frag-F | GGCTACGGTCTCTCCTTTAATTAAAATTGTTTTTTTGAATGATATGGTTG |
| p11-Frag-R | GGCTACGGTCTCTTGATCGATTGCCTATTAAC |
| p11-BB-F | GGCTACGGTCTCTATCAGATCTTGCGATGTAAATGTTGCCATGTG |
| p11-BB-R | GGCTACGGTCTCTAAGGCATCAAATAAAACGAAAG |
| p12-Frag-F | GGCTACGGTCTCAACTAGTATACTATTTCAAAACATATTTG |
| p12-Frag-R | GGCTACGGTCTCGCAACATTTACATTCCTCATTTTCCATCATC |
| p12-BB-F | GGCTACGGTCTCGGTTGCCATGTGTATGTGG |
| p12-BB-R | GGCTACGGTCTCATAGTGTCTTTCTTAATTAAAGGCATCAAATAAAACG |
| p13-Frag-F | GGCTACGGTCTCACTTAAATAATGTAATACGAGCTCG |
| p13-Frag-R | GGCTACGGTCTCGTTCCATACCTCAGGTTTGC |
| p13-BB-F | GGCTACGGTCTCGGGAATGTAAATGTTGCCATGTG |
| p13-BB-R | GGCTACGGTCTCATAAGTTACAACTTAATTAAAGGCATCAAATAAAACG |
| p14-Frag-F | GGCTACGGTCTCTTACGTAGTTTGTGTCCAGTATCC |
| p14-Frag-R | GGCTACGGTCTCCCAACATTTACATCAAAACAGGTGCAGCGGC |
| p14-BB-F | GGCTACGGTCTCCGTTGCCATGTGTATGTGG |
| p14-BB-R | GGCTACGGTCTCTCGTAAAATATGAACAATTTTAATTAAAGGCATCAAATAAAACG |
| p15-Frag-F | GGCTACGGTCTCGCCTTTAATTAATGAACTTATATCTTCTGAACTTTTTTAACGTCTTG |
| p15-Frag-R | GGCTACGGTCTCGTGGCAACATTTACATGTCGGATACCACCGAGCC |
| p15-BB-F | GGCTACGGTCTCGGCCATGTGTATGTGGGCG |
| p15-BB-R | GGCTACGGTCTCGAAGGCATCAAATAAAACGAAAGGC |
| p16-Frag-F | GGCTACGGTCTCGCAAACTTTAAATGTGCCTCG |
| p16-Frag-R | GGCTACGGTCTCGTCCAACACCGTCTTGGAATG |
| p16-BB-F | GGCTACGGTCTCGTGGATGTAAATGTTGCCATGTG |
| p16-BB-R | GGCTACGGTCTCGTTTGCCCCCGCATCTTAATTAAAGGCATCAAATAAAACG |
| p17-Frag-F | GGCTACGGTCTCAACCCTCTTTTTGGAATCCTG |
| p17-Frag-R | GGCTACGGTCTCAATCGGAAGCGAGAAAATCGC |
| p17-BB-F | GGCTACGGTCTCACGATGTAAATGTTGCCATGTG |
| p17-BB-R | GGCTACGGTCTCAGGGTATTTTAATTAAAGGCATCAAATAAAACG |
| p18-Frag-F | GGCTACGGTCTCTTCTTTTCGGCGGATTGATAG |
| p18-Frag-R | GGCTACGGTCTCTCTCCCTTGGCTAATGCGAATATAAAAC |
| p18-BB-F | GGCTACGGTCTCTGGAGATGTAAATGTTGCCATGTG |
| p18-BB-R | GGCTACGGTCTCTAAGAAATAGTTAAACCTTAATTAAAGGCATCAAATAAAACG |
| pO54-Frag-F | GGCTACGGTCTCAGCGGCTTAATTTCGCTCG |
| pO54-Frag-R | GGCTACGGTCTCTCACATGGCAACATTTACATATGTTCATGACTCCATTATTATTGTAC |
| pO54-BB-F | GGCTACGGTCTCTTGTGTATGTGGGCGTACG |
| pO54-BB-R | GGCTACGGTCTCACCGCGCTTGCGCAGATTAATTAAAGGCATCAAATAAAACGAAAG |
| p-Col-F | GTCTTTCGACTGAGCCTTTCG |
| p-Col-R | CCACATACCAAACCTTCCTTCG |
| **Construction of Mango-pSEVA2311 plasmid** | |
| Lib1-F | CCTTTAATTAAATGTAAATGTTGCCATGTGTATGTG |
| Lib2-R | TCCAAGACTAGTACGCGCTACATCCGCTTTAG |
| Lib3-F | GGATGTAGCGCGTACTAGTCTTGGACTCCTGTTG |

| | |
|---|---|
| Lib4-R | GCAACATTTACATTTAATTAAAGGCATCAAATAAAACGAAAG |
| Lib7-F | GTCCTACTCAGGAGAGCGTTC |
| Lib8-R | GCCATGAATGATCCCGAAGG |
| **Sequencing of ARES plasmid library** | |
| SeqDNA forward | ACACTCTTTCCCTACACGACGCTCTTCCGATCTN $_{5\text{-}7}$-GACGCAGGGTGTCGGATG |
| SeqDNA reverse | GTGACTGGAGTTCAGACGTGTGCTCTTCCGATCTN $_{5\text{-}7}$-TGGTGCTCGAGCTTGTACAG |
| **Sequencing of in vitro transcribed RNA** | |
| SeqRNA forward | AATGATACGGCGACCACCGAGATCTACACTCTTTCCCTACACGACGCTCTTCCGATCGAG |
| SeqRNA1 reverse | AGACGTGTGCTCTTCCGATCTCCGACCGTCTCAGATGGACC |
| SeqRNA2 reverse | CAAGCAGAAGACGGCATACGAGATGATCTGGTGACTGGAGTTCAGACGTGTGCTCTTCCGATCT |

### Sequence for validation

**Table S2:** The sequences and original IDs of the 18 randomly selected ARES, each 200 nt in length, are listed below for the validation experiment.

| ID | Original IDs | Sequence |
|---|---|---|
| p1 | ARES00610543 | gatatcccgtataacatttgtacactccgttatggtctgtaataaaatttgggatcgtttttttgcgttctaatttac<br>Agatataggctagagtaactactcgcaaacagtgaagatgtgcagtaagggtcaaccggcatagaattggc<br>tcgtgagtacgtcgatctaaagagactggtgtgcgctggtgcctgctga |
| P2 | ARES00171465 | cgactgtcttaaggttaagacaatcttattgtgaggtactggggtattggatgcagcatgggtacatttggact<br>acattatggatgggggttctgctacagtgtctacttgggacaatatttgcatcactataatgtGcaaaaagaa<br>aagtcgggcccttgtttagaaggtatggtcatgtaattgtatatgtctcgtgg |
| P3 | ARES00389971 | atttcgtgataatgagctgagttccttgataaagcagtttgtgtaatgaatgaatacatagattattaaatgga<br>attgacggctccttataacactccatatgatttgggaacggtgtatgcacaacgctgaggttGgaagtaatg<br>gtgattatgatagttaggtggaataaacggctttaaatataattttacccctg |
| P4 | ARES00141878 | tgttataattggatttattctactcttcctcataaccgggcttgggaacgaatctagcatcaaacttggcGgag<br>agatgtcgtttaggagtatgcatatatttctggaaatttcgggatttttgttaaggaatttccagtaggtgttgttt<br>cgtcaggaatgaaactatagcagctacattgtgatattctttcgggggt |
| P5 | ARES00013532 | gttgtcctgattatggtagtactagtagagacaacgtcatcctataaatcgtccaatatataggtgttttgggaa<br>ctgtcgaattgtgcgtacctatGatcgtcttttccgtaactaagacaagtcagcatattgggggtcgcatgtaa<br>cgattatcgcacagaagtgagtagtctaggtgatatgatgaggcggggcat |
| P6 | ARES00256315 | aagtgagattcattaaatattattggggaagaattggcttgcgagtggaaaaaatgactgatattaggataat<br>ctttttttgggatcaggtttagcatcagagttagatGagggcaagcaggtgttgattggaggtgcggaacagta<br>ctataactaccatgcgctgcttgtgggttttagacatcggatacggtggtgtcc |
| P7 | ARES00413161 | taagcgagattcaagaatttttagctttatcgtcgtggtgctgtcatgtagtaagtattgtgcaccgggcttagt<br>ctgttgggagtcggggtagttaaagagttgtGaatgatcaaaggtatgtctatgactcagtccatagatcggg<br>aatagtgtggtcttccacctgtcactgtggaatggaattatactacacgctg |
| P8 | ARES00523631 | aggattttgaataaagtggtaaagttgttgttgctagtgtctgaagatggtatgtttcgtaacgtctgcattgcc<br>aaaataagtttgagcttatttactttcgggtgaatttgggacaataatttactggttcataaattGggggtaagt<br>agttacaatatcagaagtacgttggatgtactaatttagccatatagcgg |
| P9 | ARES00074902 | aattgttcatattttacgtagtttgtgtccagtatcccagatacagaccgattagttttgcacgatctgtaaggag<br>ggatgggattttgcatgccggtacgcttttgatttaaatGggacgaagctggtgcagcgattcttgatcaagta<br>tagggagagttagggggtatgtatgtgtactagccgctgcacctgttttg |
| P10 | ARES00116280 | gatgcgggggcaaactttaaatgtgcctcgtgaacacgcggttgaacaattgaaatgtgagaggtgaggttg<br>tttagtgtcattggaaatgattttcgattaaactgggcgttagtggatGcgtcaaatttgtaaaactaaggtgtt<br>gtcgacaatcgtgtcaggaaagctgccttgtaggcattccaagacggtgttgg |
| P11 | ARES00237232 | gaaagacactagtatactatttcaaaacatatttgtggaatgaaattgcttatgccgtgatctacagtgaagg<br>gcggcgttctttccggaaagctgttggggacaccagattgattttgtcaatttcaaatacactttaaGatggcg<br>gtactaaattataacacttaccttgctagctagcggatgatggaaaatgagga |
| P12 | ARES00287266 | agagtatgatcccattggttctatagggaatgaaacatttgttaacatacagtggtatgctgatttttgcacggt<br>ctgtgcatgtaggccgtctGtctgtaatgctgtgtgaaacaagaatcgtcgagtgcggttaataggggggcttg<br>ttttcgctttccgatcgcgagaatgttattttctatcctatcaatggtttga |
| P13 | ARES00086654 | tgaacttatatcttctgaactttttttaacgtcttgttggttaccattagattaaaatagctatagaagcggtagat<br>cgtacgttcctcctaagagatgtcaaatatgtattcattacagtggattttgatactttgttgaaagttcgaaaa<br>Ggtattaagatggttggtttagggttgttaaggctcggtggtatccgac |
| P14 | ARES00422720 | gttgtaacttaaataatgtaatacgagctcggtagtatctttagcagtatttatcacttggttggctaaattccat<br>aatgcgaaagaacagggggggaacatgattttttttccaggaggtcacgccaagattgtaaGagggctgaga<br>gagtcggaaagtgtatacatttaaaggatgtccccgcaaacctgaggtatgga |
| P15 | ARES00952673 | ggtttaactatttcttttcggcggattgatagtttaacatctcgtgtgttgatggccaaggtgttagaaggatcgt<br>gtcttgagggccgtggattgttctgatgattttggcaggaaagttaacaaaaagcaaGgatcttatagtatg<br>cattatgcttatggttttatattcgcattagccaagggag |

| | | |
|---|---|---|
| P16 | ARES00173427 | aataccctctttttggaatcctgccatgatcggactttgtctaggtgggagctttttcaaacgattttagagtaatgatgacgtactttttgtatGggctcaattgtgtgtttaggacctttcagccatagtattctcaggctattacagacgagtacacttcattgcataactgtaaagggcgattttctcgcttccg |
| P17 | ARES00432513 | aattgtttttttgaatgatatggttgcattcatatttacatttacatccatatttgtgtatagcgattttgtaatatggtagtctatgatataaatactGattgatattgccgaaacgaggtaaaaaacatttggtgagccattacatcttttgacagatttatcgaagatggttaataggcaatcgatcagatcttgcg |
| P18 | ARES00438390 | atggagtttacttcgggtaacattggttatatttgagatttgacacatgaccgcactagacttcactacgtgggtttaaacgtttactggttattttcagattttgtatgaaagttgaatatagggataggGacgctagggctggagtgccgcagcctgataaagccgatacggggatatctcttaggtggtataggtag |
| pO[54] | Pos-Control | atgagctctctgcgcaagcgcggcttaatttcgctcgctccgatcattctaaaattagaaacacattgaaaaacaataccttgaagtctgttttcagaccttggcacagccgttgcttgatgtcctgcgcaacatgtacaataataatggagtcatgaacatatgg |

### Plasmid maps:

1) N200 ARES

https://benchling.com/s/seq-wG6faFrPGndrWYAhEMqq?m=slm-wg6jqKmiUQcNzv7zXjAx

2) pHH100-dBroccoli construct

https://benchling.com/s/seq-bD8GNHJIVsiaienqUajK?m=slm-NNHWA6VcrijY2MuG6UKe

3) N200-pHH100-dBroccoli construct

https://benchling.com/s/seq-VxVk2xeLeUp684jSTSbb?m=slm-LhFMu9GY5rYQlaGJzR3s

4) Mango-pSEVA2311 construct

https://benchling.com/s/seq-CfaFXRxx76D39VVFn3LS?m=slm-Q88cA84fxmZQlkUefZ72

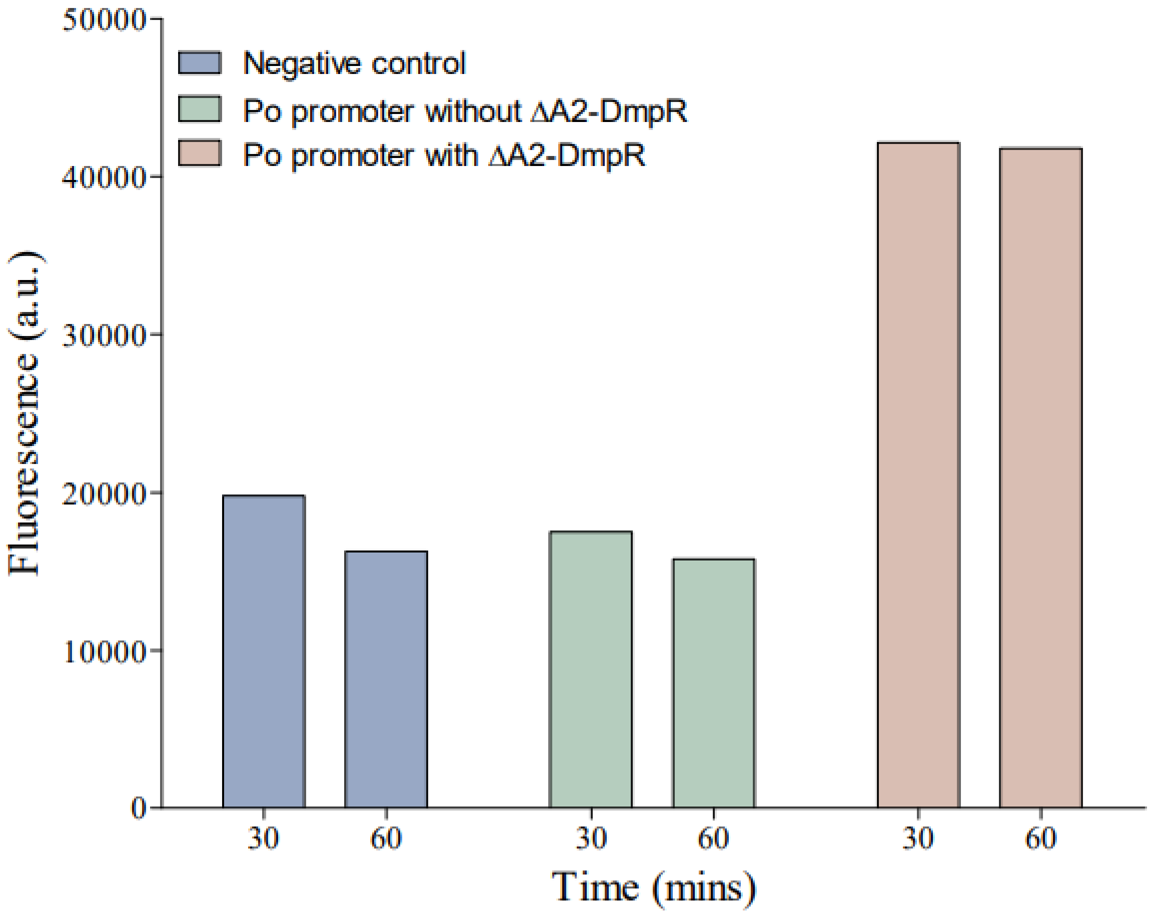

**Figure S1: Importance of ΔA2-DmpR for *P. putida* $\sigma^{54}$-Dependent IVT Reaction**

This figure illustrates the critical role of ΔA2-DmpR in the *in vitro* transcription (IVT) assays using the ARES plasmid DNA library as templates, core RNA polymerase, and $\sigma^{54}$ purified from *P. putida*.

Fluorescence of the Mango aptamer was used to measure RNA synthesis. The results are presented for three experimental groups:

1. **Negative Control:** No template added, showing baseline fluorescence.
2. **Po Promoter without ΔA2-DmpR:** Template of the Po promoter without ΔA2-DmpR, showing fluorescence levels similar to the negative control.
3. **Po Promoter with ΔA2-DmpR:** Template of the Po promoter with ΔA2-DmpR, showing significantly higher fluorescence levels compared to the negative control.

The data demonstrate that RNA synthesis does not occur in the absence of ΔA2-DmpR, as indicated by the similar fluorescence levels of the negative control and the Po promoter without ΔA2-DmpR. In contrast, significant RNA synthesis, indicated by increased fluorescence, is observed in the presence of both the Po promoter template and ΔA2-DmpR. This highlights the essential role of ΔA2-DmpR in initiating transcription from the σ54 dependent Po promoter.

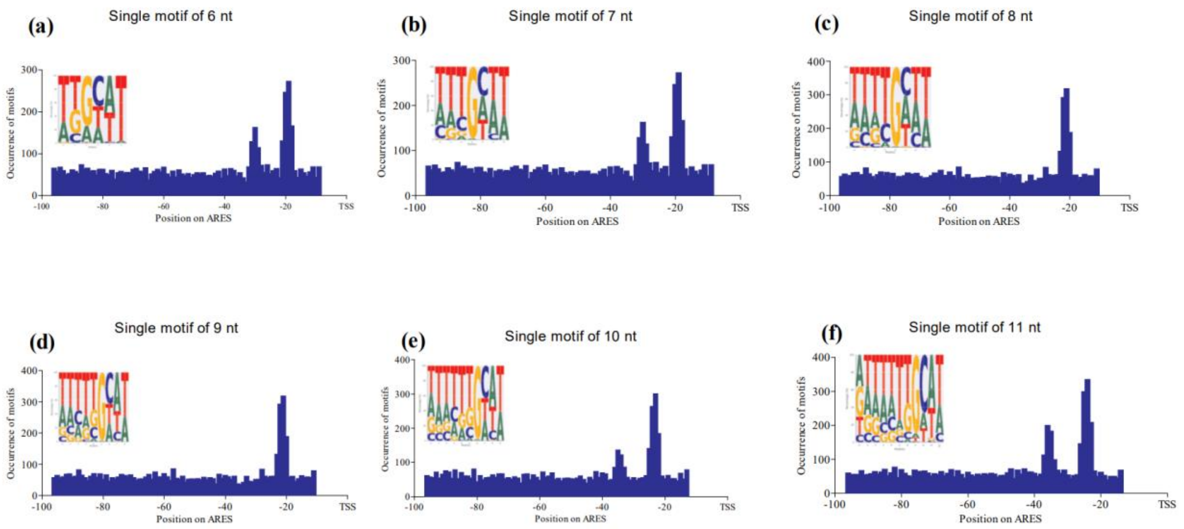

**Figure S2: Distribution of single motif within a 97 bp Region Upstream of the TSS in the Highest Expressing ARES Subset**

This figure illustrates the distribution of a single conserved motif within a 97 bp region upstream of the Transcription Start Site (TSS) across the highest expressing ARES library for motifs ranging from 6 to 11 nucleotides in length. The x-axis represents the position relative to the TSS, and the y-axis indicates the occurrence frequency of each motif.

(a) The 6 bp motif (TTGCAT) is identified at the -12 position, consistent with previous core promoter analyses.

(b) The 7 bp motif shows a sequence extension, adding a T at the beginning (TTTGCTT).

(c) The 8 bp motif extends further with an additional T (TTTTGCTT).

(d) The 9 bp motif continues this pattern (TTTTTGCAT).

(e) The 10 bp motif follows the same trend (TTTTTTGCAT).

(f) The 11 bp motif (ATTTTTTGCAT) displays the final extension in this analysis.

Each motif is consistently located at the same position upstream of the TSS, with the sequence extension demonstrating high conservation across the lengths studied.

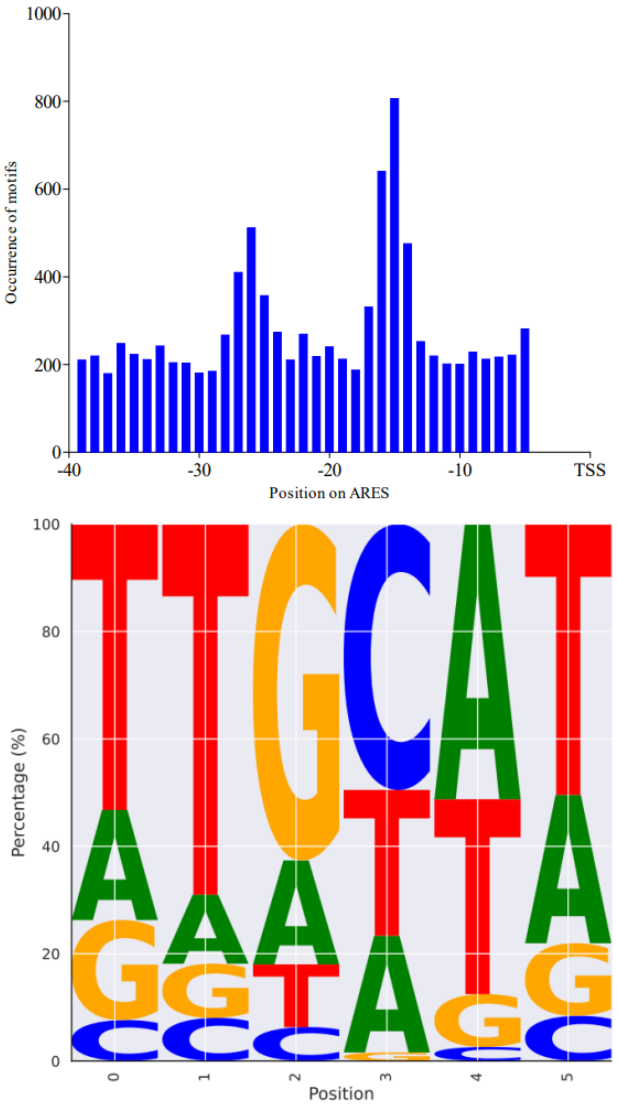

**Figure S3: Distribution of single motif within the 40 bp Region Upstream of the TSS in the Highest Expressing ARES Subset**

This figure illustrates the distribution of a single conserved motif within a 40 bp region upstream of the TSS among the highest expressing ARES subset. The x-axis represents the position relative to the TSS, and the y-axis indicates the occurrence frequency of the motif. Analysis reaffirmed the presence of the conserved 6 bp motif (TTGCAT) at the -12 position.

**Table S3:** Annotation of IVT-generated $\sigma^{54}$-RNA polymerase binding motifs in the upstream regions of genes in the *P. putida* genome.

| Gene | Product | -24 motifs | -12 motifs | p-value |
|---|---|---|---|---|
| PP_2783 | 3-oxoacyl-ACP reductase family protein | TGGGCT | TGGCAT | 0,00405 |
| PP_4970 | cytochrome c | GTGGCT | GTGTTT | 0,00353 |
| PP_1666 | DUF2066 domain-containing protein | GTGGTT | GTGTAT | 0,003 |
| eutC | ethanolamine ammonia-lyase subunit EutC | TTGGCA | TTGCAT | 0,000286 |
| PP_1786 | glycosyltransferase | ATGGCA | TTGCAT | 0,000286 |
| PP_1030 | hypothetical protein | TGGGCT | TTGATT | 0,000918 |
| PP_1810 | hypothetical protein | TTGGCT | TTGTAT | 0,00126 |
| PP_1923 | hypothetical protein | ATGGCT | TGGCAT | 0,00405 |
| PP_0091 | lipoprotein | CTGGCA | TGGCTT | 0,00462 |
| PP_1622 | peptidoglycan DD-metalloendopeptidase family protein | TTGGTT | TTGCAA | 0,00329 |
| PP_4204 | type II toxin-antitoxin system MqsA family antitoxin | TTGGCA | GTGTAT | 0,003 |
| PP_1394 | 5-guanidino-2-oxopentanoate decarboxylase | ATGGCT | GTGTTT | 0,00353 |
| PP_4577 | 5-oxoprolinase subunit PxpA | TGGGTA | GTGTTT | 0,00353 |
| PP_0859 | amidohydrolase | CTGGTA | GTGCTT | 0,0015 |
| PP_3521 | DMT family transporter | GTGGTT | TTGTAT | 0,00126 |
| PP_1210 | Dps family protein | TCGGCA | TTGAGT | 0,00381 |
| PP_1584 | DUF2514 domain-containing protein | TGGGCA | GTGCAT | 0,00174 |
| PP_3248 | Dyp-type peroxidase | TTGATA | TTGCAA | 0,00329 |
| motA | flagellar motor stator protein MotA | TTGGTT | TGGCAT | 0,00405 |
| purT | formate-dependent phosphoribosylglycinamide formyltransferase | GTGGCT | TGGCAT | 0,00405 |
| pgi | glucose-6-phosphate isomerase | TTGATA | TGGCTT | 0,00462 |
| PP_4543 | GNAT family N-acetyltransferase | TTGTTA | TTGAGT | 0,00381 |
| PP_1159 | hypothetical protein | TTGGCT | TTGCAT | 0,000286 |
| PP_5488 | hypothetical protein | ATGGCT | GTGTAT | 0,003 |
| PP_0182 | hypothetical protein | ATGGCA | TTGCAT | 0,000286 |
| PP_2952 | LysR family transcriptional regulator | GTGGTT | TGGCAT | 0,00405 |
| PP_3132 | oligosaccharide flippase family protein | TGGGCA | TAGCAT | 0,00381 |
| PP_5043 | PhoPQ-activated pathogenicity-related family protein | TTGGCT | TTGCTT | 0,000286 |
| edd | phosphogluconate dehydratase | GTGGCA | TTGTTT | 0,00208 |
| PP_4924 | S8 family serine peptidase | GTGGCA | TTGTTT | 0,00208 |
| PP_0146 | TerC family protein | CTGGTA | GTGCAT | 0,00174 |
| PP_2420 | TonB-dependent receptor | ATGGTT | TTGCGT | 0,00267 |
| yegQ | tRNA 5-hydroxyuridine modification protein YegQ | ATGGCT | GTGATT | 0,00271 |
| dusA | tRNA dihydrouridine(20/20a) synthase DusA | GTGGTA | GTGAAT | 0,00248 |
| tssG | type VI secretion system baseplate subunit TssG | TCGGCA | GTGTTT | 0,00353 |
| PP_2433 | antitoxin Xre/MbcA/ParS toxin-binding domain-containing protein | CTGGCA | TGGCTT | 0,00462 |

| | | | | |
|---|---|---|---|---|
| cls | cardiolipin synthase | TCGGCA | ATGCAT | 0,00491 |
| PP_1188 | dicarboxylate/amino acid:cation symporter | TTGGCA | TTGCTA | 0,0041 |
| PP_3989 | DNA cytosine methyltransferase | ATGGTA | TGGCAT | 0,00405 |
| PP_2810 | DUF1329 domain-containing protein | CTGGCA | TTGCGT | 0,00267 |
| PP_5363 | DUF3617 domain-containing protein | GTGGTA | ATGCAT | 0,00491 |
| flgA | flagellar basal body P-ring formation chaperone FlgA | TCGGCA | TTGCTT | 0,000286 |
| flgB | flagellar basal body rod protein FlgB | TTGGCA | TTGCTA | 0,0041 |
| flgF | flagellar basal-body rod protein FlgF | TTGGTT | TTGCTT | 0,000286 |
| flgG | flagellar basal-body rod protein FlgG | ATGGCT | TTGCAA | 0,00329 |
| fliE | flagellar hook-basal body complex protein FliE | CTGGCA | TTGCTT | 0,000286 |
| PP_2836 | fumarylacetoacetate hydrolase family protein | GTGGCT | TTGCAA | 0,00329 |
| PP_1946 | glucose 1-dehydrogenase | TTGGCA | TTGCAA | 0,00329 |
| PP_0298 | GlxA family transcriptional regulator | TAGGCA | TTGCGT | 0,00267 |
| PP_2828 | hypothetical protein | TAGGTA | GTGCAT | 0,00174 |
| PP_3592 | MurR/RpiR family transcriptional regulator | ATGGCT | TTGCAA | 0,00329 |
| ntrB | nitrogen regulation protein NR(II) | GTGGTT | TTGCAT | 0,000286 |
| PP_0952 | RNA polymerase factor sigma-54 | AAGGCA | TTGCTT | 0,000286 |
| PP_3288 | universal stress protein | TCGGTA | TTGAGT | 0,00381 |
| urtA | urea ABC transporter substrate-binding protein | TAGGCA | TTGCAA | 0,00329 |
| PP_2209 | 2-aminoethylphosphonate--pyruvate transaminase | ATGGCA | TTGCAA | 0,00329 |
| PP_3486 | ABC transporter substrate-binding protein | ATGGCT | GTGCTT | 0,0015 |
| PP_1141 | branched-chain amino acid ABC transporter substrate-binding protein | TGGGCT | TGGCAT | 0,00405 |
| bcsB | cellulose biosynthesis cyclic di-GMP-binding regulatory protein BcsB | GTGGTT | TTGCGT | 0,00267 |
| ppx | exopolyphosphatase | ATGGCA | TTGCAT | 0,000286 |
| PP_3703 | hypothetical protein | TGGGCA | GTGTAT | 0,003 |
| PP_0052 | MBL fold metallo-hydrolase | GTGGCA | TGGCAT | 0,00405 |
| PP_3539 | MerR family DNA-binding transcriptional regulator | GTGGTT | TTGAGT | 0,00381 |
| PP_3060 | phage major tail tube protein | GTGGTT | GTGCTT | 0,0015 |
| PP_3778 | pyrroline-5-carboxylate reductase | TTGACA | TTGCTA | 0,0041 |
| PP_1450 | HlpA activation/secretion protein HlpB | GTGGCT | TAGCAT | 0,00381 |
| PP_5722 | type II toxin-antitoxin system PemK/MazF family toxin | CTGGTA | TTGAAT | 0,000631 |
| xdhA | xanthine dehydrogenase small subunit | TGGGCA | GTGCAT | 0,00174 |
